# Introducing Novel Molecular-based Method for Quantification of Homologous Recombination Efficiency

**DOI:** 10.1101/2021.01.17.427032

**Authors:** Mustapha Dibbasey, Terry Gaymes

## Abstract

**Background:** Homologous recombination (HR) pathway is a DNA double-stranded breaks repair pathway well-known for its high level of accuracy. Low HR pathway efficiency clinically known as homologous recombination deficiency (HRD) was identified in some cancers such as breast and ovarian cancers and studies have reported the sensitivity of HRD cancer cells to DNA repair inhibitors such as Olaparib. However, current techniques including immunofluorescence-based technique are qualitative-based, hence lack sensitivity to determine the functionality of HR pathway. Additionally, some of the techniques including gene expression arrays require expression study of wide range genes involve in HR pathway, which is not cost-effective. The aim of the study is to optimise a PCR-based assay (Norgen’s Homologous Recombination kit) that can be employed to quantitate HR efficiency in cells, which accurately reflects the functional status of HR pathway.

**Methods and Findings:** The kit has two test plasmids (dl-1 and dl-2) with partial deletions in the LacZ gene and the plasmids are generated from modification of pUC19. HR-proficient (HeLa and AsPC-1) and HR-deficient (CAPAN-1 cells) cancer cell lines were transfected with the two plasmids to generate functional LacZ gene (i.e. recombinant product). The recombinant product was quantified by real-time PCR. Although recombinant product was generated in all the cell lines, our real-time PCR demonstrated a high quantity of recombinant product in HeLa cell line whilst low quantity in CAPAN-1 and AsPC-1 cell lines. The quantity of recombinant product generated and quantified reflects HR pathway efficiency.

**Conclusion:** Overall, the results have provided some evidence that the PCR-based kit can be suitably employed for quantification of HR efficiency provided appropriate transfection method and reagent are used. However, further study is required to confirm HR efficiency status of AsPC-1 cells to ascertain the low HR efficiency detected by the kit in these cells.

## INTRODUCTION

The capacity to accurately repair double-stranded breaks (DSBs) is central to the normal cell growth and function. DSBs is a major underlying cause genomic instability following inaccurate repair. Hence this toxic lesion must be accurately repaired in its entirety to preserve the integrity of the genome and avert the induction of abnormal cellular proliferation and growth, which gives rise to cancer cells (Hanahan and Weinberg 2011; Georgoulis et al. 2017). In addition to exogenous insults such as UV light, replication process is another source of DSB. The attempt to replicate over an unrepaired SSB in the S phase by replication machinery induces stalled replication fork and DSB, which requires DSB repair pathways as demonstrated both in the yeast and mammalian cells to initiate the repair process and restore normality (Arcangeoli and de Lahondes 2000; Jasin and Rothstein 2012; Mehta and Haber 2014; Shimizu *et al*. 2017). The two most important pathways associated with DNA DSBs repair are classical non-homologous enjoining (NHEJ) and homologous recombination (HR) that play indispensable role in ensuring normal cellular growth and function as well as cell cycle progression (Rodriquez and D’Andrea 2017). Unlike NHEJ, HR repair pathway is active only in late S and G2 phases in mitosis, during which sister chromatids are available. HR repair pathway is associated with high degree of fidelity (i.e. accurate DNA DSB repair) owing to the presence of sister chromatid as a template to guide the repair process, hence labelled as error-free pathway which is important for genomic stability (Frey and Pothuri 2017). HR pathway is intricately linked to another essential DSB repair pathway known as Fanconi pathway to repair interstrand/intrastrand crosslinking of DNA alpha-helix structure during DNA synthesis which also contributes to ensuring genomic stability. For instance, the defect in the Fanconi pathway is a fatal consequence of Fanconi anaemia, a fatal disorder associated with wide spectrum of clinical phenotypes and bone marrow failure (Rodríguez and D’Andrea 2017; Nalepa and Clapp 2018). HR is also essential during meiosis (specifically at prophase l) involving sperm cells and ova leading to new combination of genetic sequences via genetic recombination. This new combination of genetic sequences elucidates the greater variations at molecular level among offspring, which is important for the population to adapt during evolution (Lam and Keeney 2014; Szekvolgyi et al. 2015). To further expound its biological significance and universality, HR mechanism also exists in bacteria and virus for the horizontal gene transfer, in which genetic materials is being exchanged between different strains (Gonzalez-Torres et al. 2019).

As HR repair mechanism is tasked to ensure accurate DSBs repair and genomic integrity in S and G2 phases of the cell cycle, its deficiency suggests a fatal consequence. HR deficiency (HRD) is clinically defined as functional inactivation of HR repair mechanism in the cells due genetic or epigenetic inactivation of HR repair genes such as BRCA1 and BRCA2 genes. In fact, HRD is one the underlying causes of genomic instability, an enabling characteristic feature of cancer cells (Hanahan and Weinberg 2011). HRD was first identified in breast cancer largely as a consequent of germline mutation in BRCA1/2 genes. Also, Cancer Genome Atlas Study analysis has revealed that around 50% of ovarian cancer cases have HRD which were mainly due to germline mutations in BRCA1/2 genes, accounting for 14 to 15% of ovarian cancer cases (The Cancer Genome Atlas Research Network, 2011; Frey and Pothuri et al. 2017). The lifetime risk of developing breast cancer at the age of 80 is estimated to be around 72% and 69% for BRCA1 and BRCA2 carrier women, respectively. On the contrary, it is estimated that around 44% and 17% of women with germline mutation in BRCA1 and BRCA2 genes, respectively, will develop ovarian cancer at the age of 80 (Kuchenbaecker et al. 2017). Loss-of-function mutation in other HR repair genes such as ATM, Rad50 and Rad51 are also implicated to have association with increased risk of developing breast and ovarian cancer (Cerbinskaite et al. 2012). HRD was also observed in other malignant solid tumours including endometrial (34.4%), bladder (23.9%), malignant melanoma (18.1%), pancreatic (15.4%) and colorectal (15.0%). The HRD arises as an aftermath of loss-of-functional mutations (germline or somatic) in HR genes (e.g. BRCA1/2, ATM, RAD50, RAD51, CtlP and MRE11) that modulate HR repair mechanism in cells (Cerbinskaite et al. 2012). The most mutated HR genes included BRCA2 (3.0%), BRCA1 (2.8%), ATM (1.3%), and ATR (1.3%). The mutations that were found in these HR genes were frameshift insertions or deletions whilst point mutations such as missense and nonsense mutation were rarely found. Hypermethylation of promoter regions was also found in BRCA1 gene leading to epigenetic inactivation (Heeke et al. 2018). Beyond solid malignant tumours, HRD is also implicated in haematological malignancies especially acute myeloid leukaemia (AML) (Gaymes et al. 2002; Gaymes et al. 2013; Cream et al. 2017). Another recent study of Gaymes et al. (2013) identified mono-allelic mutations in CtlP and MRE11 from primary AML cells and cell lines which lead to downregulation of HR repair mechanisms. Evidence from the previous study on AML has demonstrated overactive NHEJ pathway in mammalian cells, an attributable feature that arises largely due HRD in the S and G2 phases, paving away for cells to utilize error-prone MMEJ (also known as alt-NHEJ) resulting in mutagenic events in cells (Gaymes et al. 2002). The aberration of HR repair mechanism is also identified in other haematological malignancies such as B-cell lymphocytic leukaemia and lymphoma, and Fanconi anaemia (Cerbinskaite et al. 2012; Katsuki and Takata 2016).

The assays to study HRD in cells include gene expression arrays, immunofluorescence-based and immunohistochemistry-based techniques to determine focus formation (yH2AX/RAD51 foci), methylation-specific microarrays, and direct Repeat-Green Fluorescence Protein (DR-GFP) (Ohashi et al. 2005; Gaymes et al. 2006; Gaymes et al. 2009; Cerbinskaite et al. 2012). The current methods including the immunofluorescent-based and immunohistochemistry-based techniques are basically qualitative assays. These qualitative assays have some degree of questionable sensitivity to determine HR function in the cells since their core principle is based on localisation of HR proteins instead of holistic functional assessment of HR pathway efficiency in cells. Some of the current assays such as gene expression arrays require the assessment of many HR genes. To summarise, comprehensive evaluations of HRD in cells based on the current techniques are partly limited by the lack of uniformity in approach, sensitivity of qualitative assays and cost-effectiveness of assays for testing and defining HRD, as well as labour-intensive nature of the techniques.

Having underscored the therapeutic significance of identifying HRD in cancer patients, a rapid and sensitive quantitative-based technique that can comprehensively assess HR efficiency (i.e. the functional status of HR repair pathway) would improve both clinical and research practice. In the recent study of Mio et al. (2018), an HR kit known as Norgen’s HR kit was employed to assess the efficiency of HR pathway in a cell line treated with Bromodomain and Extra-terminal protein inhibitors. The kit was able to quantify HR efficiency in the cells exposed to the inhibitor. Also, the suitability and high throughput of the kit was well acknowledged in several other research studies (Ohba et al. 2014; Misic et al. 2016, Na et al. 2016). This kit is molecular-based technique which could provide rapid and sensitive opportunity to determine the level of HR efficiency in both bacteria and eukaryotic cells and eliminate the need for labour-intensive screening by using traditional protein expression and staining techniques (e.g. immunofluorescence-based and immunohistochemistry-based techniques). Furthermore, the molecular-based kit is also quantitative-based (real-time PCR); which means it could provide relative expression level of HR that could be translated as HR pathway efficiency (Norgen Biotek Corporation, 2010). Hence, such characteristics merit the approach to optimise this kit for further studies on HR pathway. Thus, the study aims to optimise Norgen’s HR kit that can be employed to quantitate HR efficiency in cancer cells using a sensitive molecular-based approach, real-time PCR.

## MATERIALS AND METHODS

### Cell Lines

Cervical carcinoma cell line (Hela cells), human pancreatic cancer cell line (AsPC-1 (ATCC® CRL-1682™)), Human Pancreatic Adenocarcinoma Cell Line (Capan-1, ATCC HTB-79), were provided by Dr Terry Gaymes. The cell lines were cultured and maintained in Dulbecco’s Modified Eagle Media (DMEM) (Sigma Aldrich, United Kingdom) supplemented with 10% heat-inactivated foetal bovine serum and 1% Penicillin plus streptomycin and incubated in 37^°^ C incubator with 5% CO_2_ atmosphere. The cell lines were passaged when 80% of confluence was achieved to maintain cellular growth and proliferation through-out the study period.

### Trypsinisation and Cell Counting

The prepared DMEM, Dulbecco’s Phosphate Buffered Saline (PBS) (Modified, without calcium chloride and magnesium chloride, liquid, sterile-filtered, suitable for cell culture) (Sigma Aldrich, UK), and trypsin (prepared by Lauren, laboratory manager) were placed in the water bath at 37^°^ C for 20 to 30 minutes to attain room temperature. The DMEM medium in the T75 cell culture flask (Thermo Fisher, UK) with cultured cell lines was aliquoted into 50ml falcon tube. The 10ml of modified PBS was aliquoted into the flask to remove the residual cells and neutralise acidic environment prior to the addition of trypsin-EDTA (Sigma-Aldrich, UK) which was followed by 5minutes incubation to induce detachment of the adherent cells from the surface of the T75 culture flask. After the 5minutes incubation period at 37^0^C in 5% CO_2_ atmosphere in which cells were completely detached, the trypsin-EDTA was then aliquoted from the flask and added to the 50ml falcon tube containing the DMEM medium and PBS. The falcon tube was centrifuged at 2000rpm for 5minutes and the supernatant was discarded to get pellets of cells. The cells were suspended into 20ml of fresh DMEM medium for cell counting. The 50ul of DMEM medium containing the suspended cells and 50ul of trypan blue dye (Sigma-Aldrich, UK) were aliquoted into 1.5ml microcentrifuge tubes to achieve a 1:1 ratio. Trypan blue was unable to penetrate the cellular membrane of live cells, which were counted as live cells. The live cells were counted using 4×4 FAST-READ 102® grid (Biosigma, Italy) as per manufacturer’s instruction (Biosigma S.r.l., Via Valletta, Cana, Italy) as shown in figure 7. Two counting were taken and means obtained to enhance the accuracy of obtaining 1×10^6^ cells. *The formula and calculation procedure are found in appendix 1*.

### Plasmid Transfection

The cell lines were transfected using Amaxa nucleofection system™ (Amaxa, Koeln, German), an efficient electroporation method of transferring nucleic acid materials such as DNA and RNA, and plasmids into cells especially difficult-to-transfect cell lines such as Natural Killer cells (Maasho et al. 2004). Approximately, 1×10^6^ live cells were used for transfection through-out the study period for both test and positive control transfections. Following cell counting, the required volume containing 1×10^6^ cells was aliquoted into two new 20ml falcon tubes and centrifuged to obtain the pellets of cells. The pellet of HeLa cells was suspended in 80ul of *Trans*IT-LT1 (Mirus Bio, Madison, US) transfection reagent. However, AsPC-1 and CAPAN-1 pellets were suspended in Lonza transfection reagent (SE Cell Line 4D-Nucleofector™ X Kit L, Lonza, Germany) when the Mirus transfection reagent was finished. Briefly, test and positive control transfection lines were set up as HR plasmids (dl-1 and dl-2) and positive control plasmid, respectively. Both HR plasmids (dl-1 and dl-2) and positive control plasmid were supplied with the Norgen’s Homologous Recombination kit (Norgen Biotek Corp., Thorold, ON, Canada). For each transfection, 10ul of each plasmid dl-1 and dl-2 were mixed and then added onto the prepared transfection cell line annotated test sample. The 10ul of positive plasmid was added to the positive control sample. The test and positive control samples were transferred into the provided 4D nucleofector single cuvette and nucleofected with an Amaxa Nucleofector apparatus (Amaxa). The cells were immediately transferred into a 6-well plate containing 2.5ml of fresh DMEM media and incubated for 48hours in a 37^°^ C incubator with 5% CO_2_ atmosphere.

### DNA Isolation

Following 48hours of post-transfection incubation, the total DNA from the transfected cell lines were isolated using PureLink Genomic DNA Mini kit (Invitrogen), as shown in appendix 2. Briefly, lysates were prepared by adding the proteinase K and digestion buffer (lysis buffer). The lysates were added to PureLink Spin column for the binding of genomic DNA to the column, which was then washed twice with both wash buffer 1 and 2 to purify the DNA material. The purified genomic DNA materials were later eluted using optimised 50ul PureLink Genomic Elution buffer. The eluates were used for both PCR and real-time PCR downstream applications.

### PCR and Real-time PCR

Three stages PCR (denaturation, annealing and extension) procedure recommended by the Norgen Biotek Company were employed in the study for the gel-based PCR and real-time PCR. The first phase of the study involves the transfection of HeLa cell lines. As per manufacturer’s recommendation, PCR cycling temperature and time set up are as follows; Initial denaturation at 95°C for 3 minutes then 95°C for 15 sec, annealing at 61°C for 15 sec, and extension at 72°C for 15 sec, and repeated for 35 cycles then a final extension at 72°C for 5 minutes. In the second phase of the study involving subsequent transfection of AsPC-1 and CAPAN-1 cell lines, the annealing temperature was reduced from 61^0^ C to 58^0^ C to enhance the visibility of the recombinant bands. The Assay and Universal primers supplied with Norgen’s HR kit (Norgen’s Biotek Corp., Canada) were used during both PCR and RT-PCR reaction set up. For assay and universal primer reactions, 2ul of the primers based on manufacturer’s instruction, 1ul of DNA eluate isolated from the transfected samples, and 10ul of DreamTag PCR Master Mix (2X) (MgCl2, dNTPs, Dream Tag buffer and DNA polymerase) (Thermo Fisher) were used for 20ul PCR reaction on both the test sample and positive control. Following PCR reaction, 1% (0.5g) of agarose gel was dissolved in 50ml of Tris-acetate-EDTA buffer and stained with 5ul of gel red. Gel electrophoresis was performed and then image obtained to view the PCR product bands.

Following successful recombination, real-time PCR was performed using Techne real-time PCR equipment (Cole-Parmer Instrument Company LTD, UK) to quantify the recombination product in both test and control samples. As in the case of gel-based PCR, similar temperature and time set ups repeated for 40 cycles were employed and then melt curve analysis was run during the reaction by relative fluorescence assessment at 0.5°C increments with a 10 second duration, starting at 55°C and continuing for 80 cycles. Although the same volume for primers (2ul) were maintained in the RT-PCR reaction set up, SYBR green DreamTag (10ul) was employed in the place of DreamTag. Besides, similar volume of DNA sample (1ul) was used, and the final volume was adjusted to 20ul PCR and RT-PCR reaction by Nuclease-free water.

### Ethical Approval

The study involves transfection of already available human cell lines, hence ethical approval was not required for this study.

## Statistical Analysis

Although the study does not require comprehensive statistical analysis, the gel electrophoresis images were clearly annotated, and the recombinant bands identified across three transfected cell lines. The real time PCR was employed to quantify the recombinant product. The interpretation of real-time PCR result is based on quantitation cycle (Cq). Cq (also known as Cycle threshold (Ct)) is defined as the number of cycles required for SYBR green fluorescence to be detected which is inversely proportional to the quantity of initial target sequence in the sample (i.e. the lower the Cq value the higher the quantity of target sequence in the sample). This means that the cycle at which SYBR green fluorescence intersects the threshold line is recorded as Cq value by real-time PCR analyser, as shown in real-time PCR result in the product information sheet of the kit (fig. 8) (Caraguel et al. 2011). To obtain delta Ct value, the Cq value of assay primer reaction was subtracted from universal primer reaction for both test and positive control sample. The delta Ct is further converted using the comparative Ct method formula (2^−ΔΔCt^) to calculate HR expression level of the transfected cell lines in Microsoft word excel, which is presented in bar graph.

## RESULTS

### No effects of transfection on cell morphology and viability

HeLa, AsPC-1 and CAPAN-1 cell lines were transfected with the test plasmids (plasmid dl-and dl-2) and control plasmid using AMAXA nucleofection system and incubated for 48hours at 37° C in 5% CO_2_ atmosphere. After 48hours of post-transfection incubation, the micrographs of the transfected cells were captured with FLoid™ Cell Imaging Station (40X objective) using the transparent light to assess the transfection toxicity on cellular morphology and growth pattern (Fig.9A-C). No morphological changes were observed across all the three mammalian cell lines, and the micrographs showed that around 90% of transfected cells were adherent to T75 culture flask surface based on rough estimation of adherent cells against floating cells (Fig.9A-C).

**Figure 9:**
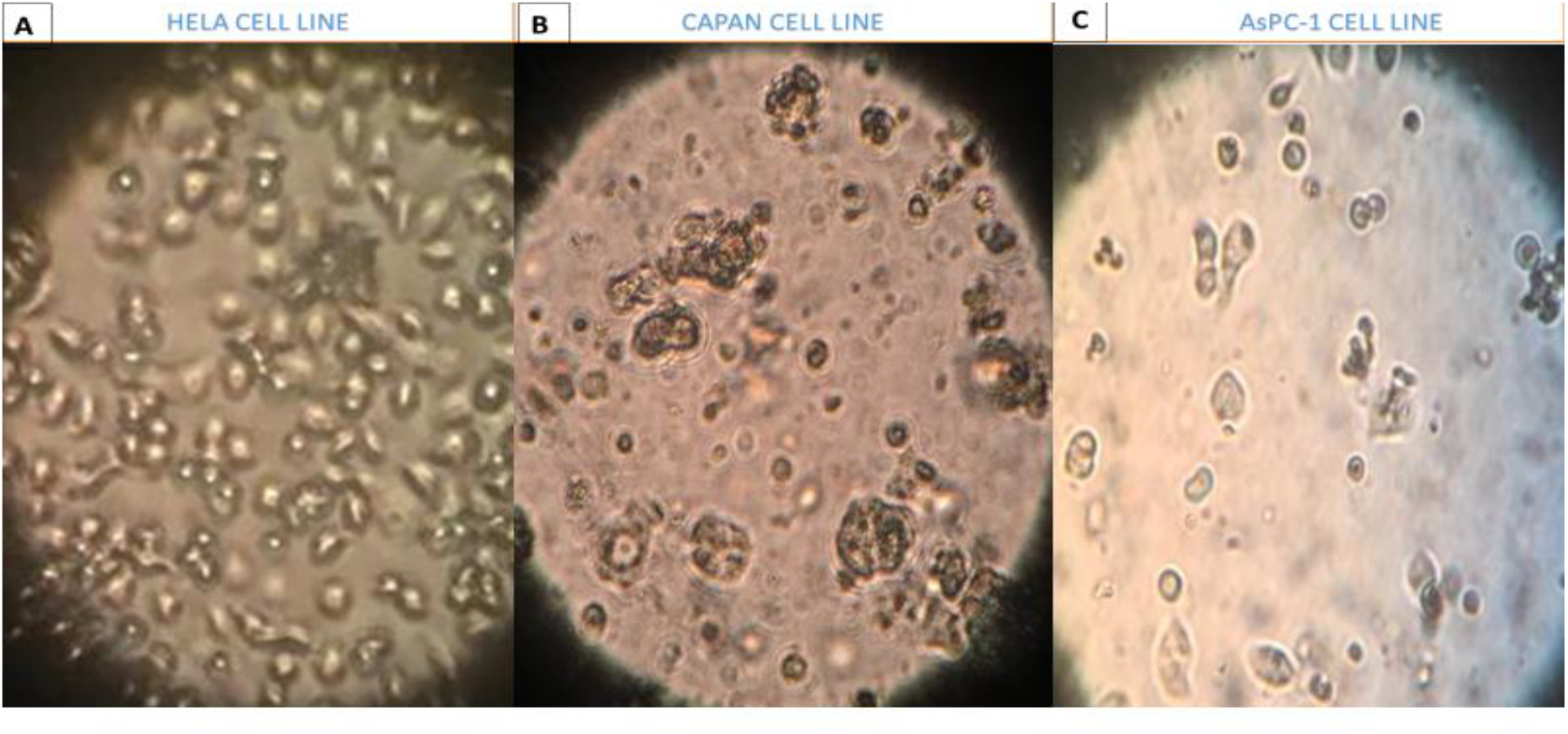
The micrograph obtained following 48hours of transfection showed monolayers of HeLa, AsPC-1 and CAPAN-1 cell lines (40X objective). **A)** The micrograph of the Hela cells showed epithelial-like cells that are attached to the surface of the flask and growing in tight patches. **B)** The micrograph of CAPAN-1 Cell line following 48hours of transfection showed cells that are round with some level of variation in size and attached to the surface of the flask in discrete patches as well as in clusters. Few apoptotic cells seen **C)** The micrograph of AsPC-1 cell line following 48hours of transfection showed epithelial-like cells that are polygonal in shape with regular dimension and variation in size as well.

### Gel Electrophoresis Results following Plasmid (dl-1 and dl-2) transfections of cell lines revealed the presence of recombinant product in all cell lines in HeLa cells, AsPC-1 and CAPAN cells

Based on the gel electrophoresis image of the transfected HeLa cell line shown in figure 10 below, recombinant product band with an actual size of 420bp was visible in the test sample with assay primer (test (A)) but not in positive control with assay primer (positive control (A)). Based on the gel image of test sample in which universal primer was employed (test (U)), two expected bands of 546bp for dl-1 plasmid backbone and 183bp for dl-2 plasmid backbone were visible. Besides, one expected band of 563bp was visible in the gel image of positive control with universal primer (positive control (U)). The 563bp amplicon commensurate with positive control plasmid amplificon size.

**Figure 10:**
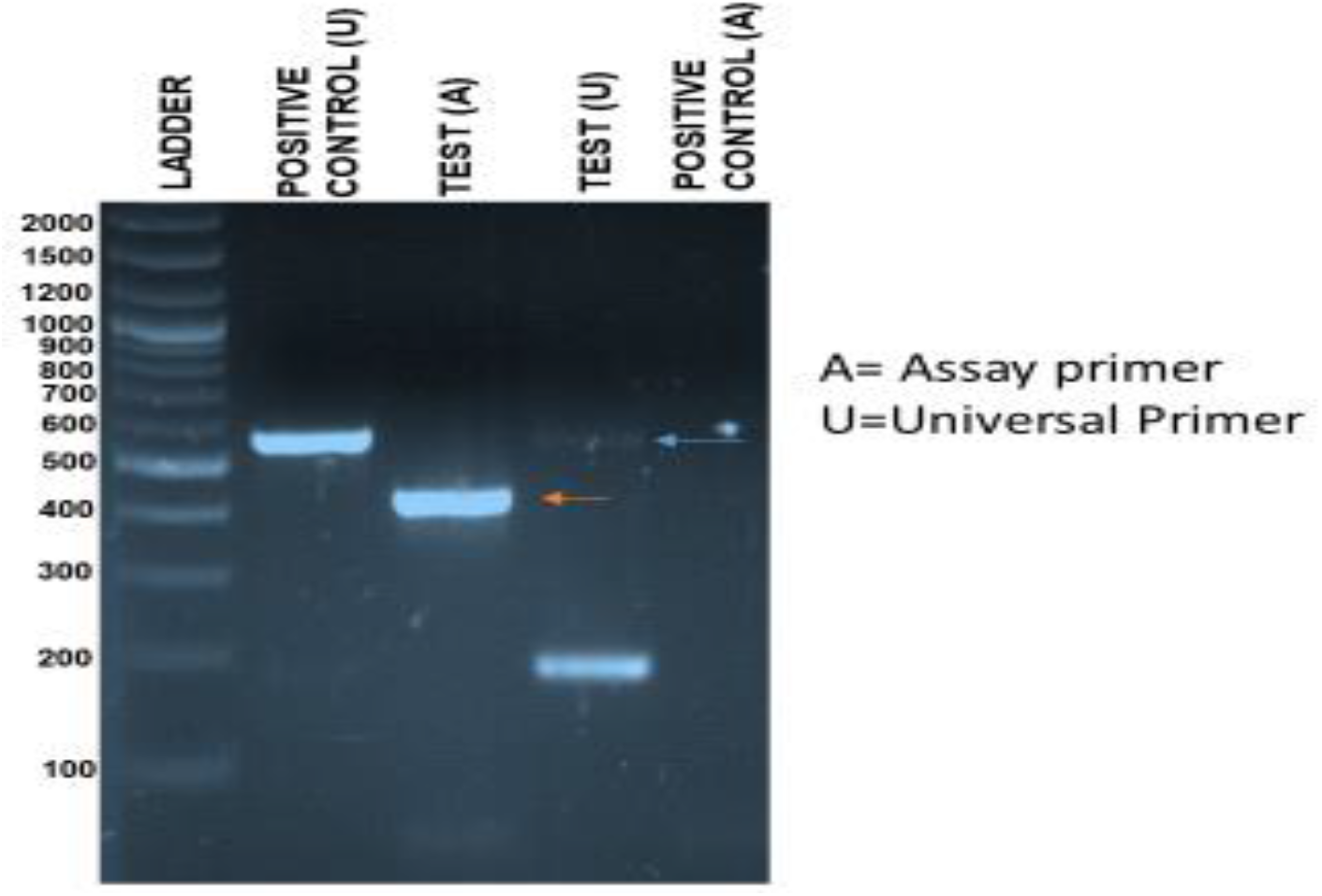
The transfection of HeLa cell line. The orange arrow indicates the presence of recombinant band (420bp) detected in the Test (A) with assay primer. The blue arrow indicates the presence of faint band. No recombinant band was visible in the positive control (A). (Test (U)= test sample with universal primer; Test (A)= test sample with assay primer; positive control (U)= positive control with universal primer; positive control (A)= positive control with assay primer; ladder=1kb plus)

### 3.2.2 AsPC-1 Cells

The gel electrophoresis image results (Fig.11A and B) consistently showed the presence of recombinant product (420bp) in the gel image of first and second transfections (A and B) of test 2(A) when assay primer was used. Expectedly, a band was visible in positive control 2(A) in both first and second transfections gel image (A and B), which commensurate with the recombinant product size of positive control plasmid (420bp). However, in the first transfection, no band was visible in the gel image (A) of both test 1(U) and positive control 1(U) in universal primer reaction. This is indicative of universal primer failure to detect and amplify both plasmid dl-1 and dl-2 backbones as well as positive control plasmid.

**Figure 11:**
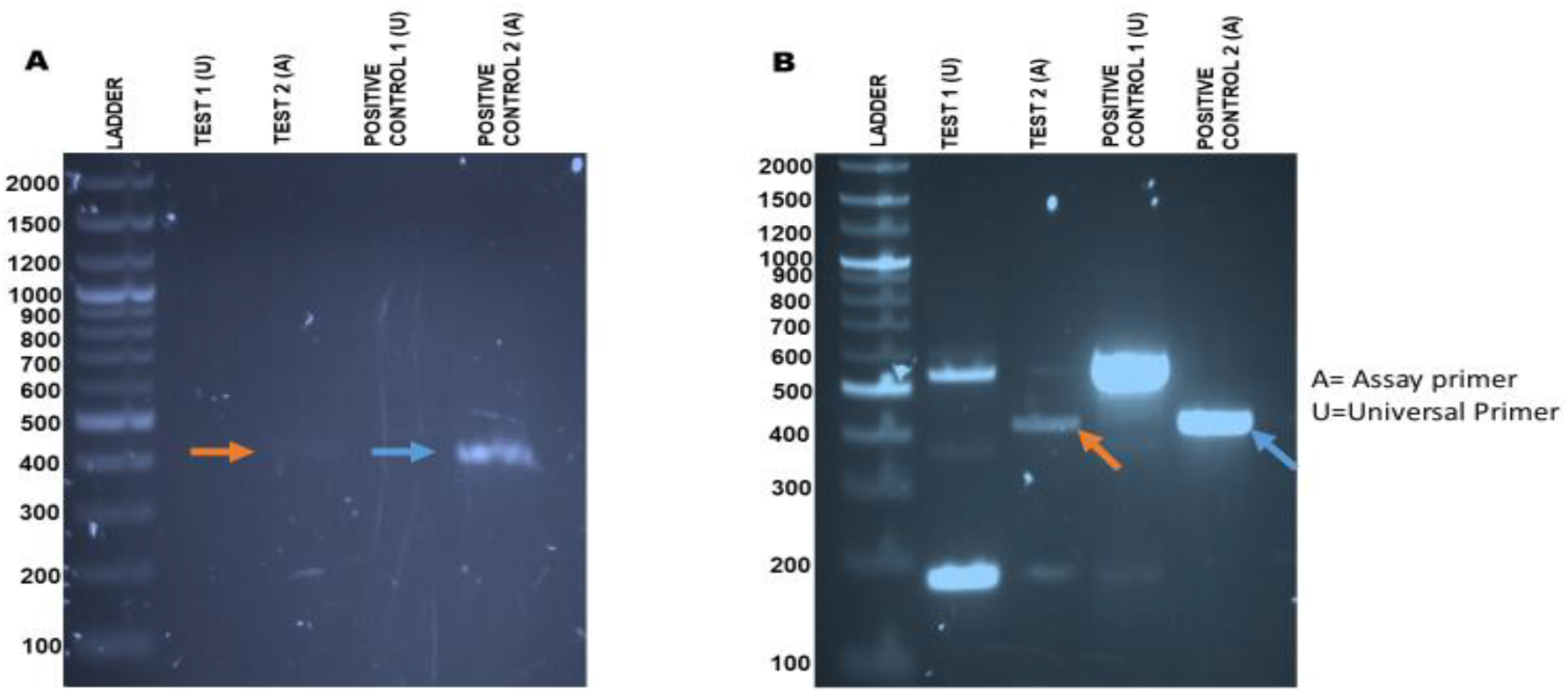
First and second transfection gel image results of AsPC-1 cell line. A and B) the pink arrow indicates the presence of homologous recombinant band (420bp) from test sample 2 and the blue arrow indicates homologous recombinant band (420bp) from positive control 2 in both first and second transfections using assay primer. (Test 1(U) = test sample with universal primer; Test 2(A)= test sample 2 with assay primer; positive control 1(U)=positive control with universal primer; positive control 2(A)= positive control with assay primer; A= First transfection gel image; B= second transfection gel image; ladder=1kb plus)

In the second transfection, several expected and non-expected (i.e. non-specific) bands were visible in the gel image (B) including the additional non-specific band of 183bp in test (2A). Based on gel image result of the test 1(U), two expected bands (546bp and 183bp) were observed owing to amplification of the backbones of both plasmids dl-1 and dl-2 by universal primer. Additionally, another non-specific band with an estimated size of 380bp was unexpectedly observed. The gel image of the positive control 1(U) following second transfection demonstrated three expected amplicons 563bp, 546bp, and 183bp that commensurate with amplicon size of positive control plasmid, plasmid dl-1 and plasmid dl-2, respectively, using the universal primer.

### CAPAN-1 Cell Line

HR deficient cell line (CAPAN-1 cell line) was established as negative control. Like HeLa cells and AsPC-1, the recombinant products (420bp) in both test 2(A) and positive control 2(A) of the first and second transfections were visible. However, primer-dimers were also observed in the test 2(A) and positive control 2(A) of the second transfection (B).

In the case of gel image result of test 1(U) of first transfection (A), two expected bands (546bp and 183bp) were visible owing to amplification of the backbone of both plasmids dl-1 and dl-2 by universal primer. Additionally, another non-specific band with an estimated size of 380bp was unexpectedly observed. For the second transfection (B), only 546bp band was visible in the gel image result of test 1(U) when universal primer was used. This band commensurate with amplicon size of plasmid dl-1 and indicates that the universal primer was able to detect and amplify only the plasmid dl-1.

The gel image result of the positive control 1(U) following gel electrophoresis of first transfection demonstrated only one visible band (563bp) when universal primer was employed. This band is genuine band as it commensurate with amplicon size of positive control plasmid. However, two bands 563bp and 183bp that commensurate with amplicon size of positive control plasmid and plasmid dl-2, respectively, were visible in the second transfection gel image (B) of positive control 1(U). Another non-specific band with an estimated amplicon size of 380bp was also detected in positive control 1(U).

### Norgen PCR-based HR kit reveals elevated HR efficiency in HeLa cells but low in AsPC-1 and CAPAN-1 cell lines

The primary aim of the study is to quantitatively measure HR efficiency in cell lines using Norgen’s HR kit, a PCR-based method of quantifying HR efficiency. Real-time PCR was subsequently performed on the extracted DNA sample from transfected cells based upon the protocol recommended by the manufacturer, Norgen Biotek Corporation, to quantitate the recombinant product. Universal primers reaction was used alongside assay primer reaction purposely to normalize the real-time PCR to determine the difference between quantity of recombinant product and, plasmid backbones (i.e. plasmid dl-1 and dl-2, as well as positive control). Cq data was generated from the amplification plot of real-time PCR results for both universal and assay primer reactions as shown in figure 13 below to determine the HR expression level, which reflects level of HR repair pathway efficiency.

**Figure 13:**
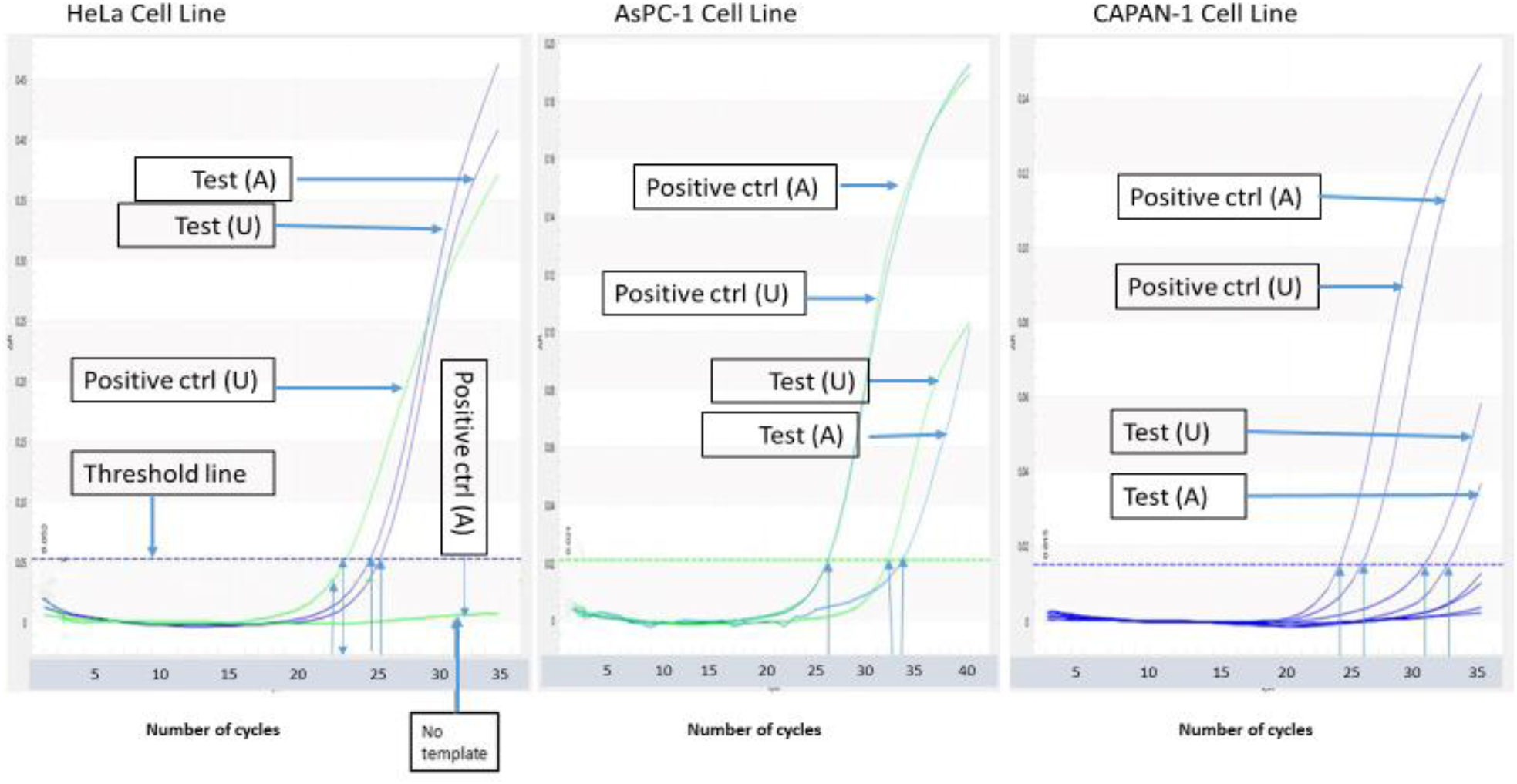
Amplification plot graph of real-time PCR results for the three cell lines. The cycle at which the amplification curve intersects the threshold line is recorded as Cq value shown by the arrow. For HeLa cells, the amplification curve of the recombinant product (test A) intersected the threshold line around cycle 25 (<30 cycle). For both AsPC-1 and CAPAN-1, the amplification curve of recombinant product (test A) intersected the threshold line beyond cycle 30 (>30 cycle). (Test (U)= test sample with universal primer; Test (A) = test sample with assay primer; Positive ctrl (U)=positive control with universal primer; Positive ctrl (A): positive control with assay primers)

For the HeLa cell line, the detection of fluorescent signal is based on the corresponding cycle number at which the threshold line is intersected by the amplification curve, which was recorded in a form of Cq value. Since Cq value is inversely proportion to the quantity of recombinant product, the amplification plot result showed a high quantity of recombinant products as the recombinant product (test A) signal was detected between cycle 20 to 25, as shown in figure 13 below, indicating a very efficient HR repair pathway (Norgen Biotek Corporation 2010; Caraguel et al. 2011). This is further justified by the Cq value analysis data shown in figure 14 in which HR expression is found to be elevated in HeLa cell line by 16-fold expression. However, no signal was detected for assay primer reaction with positive control, indicating the absence of template.

**Figure 14:**
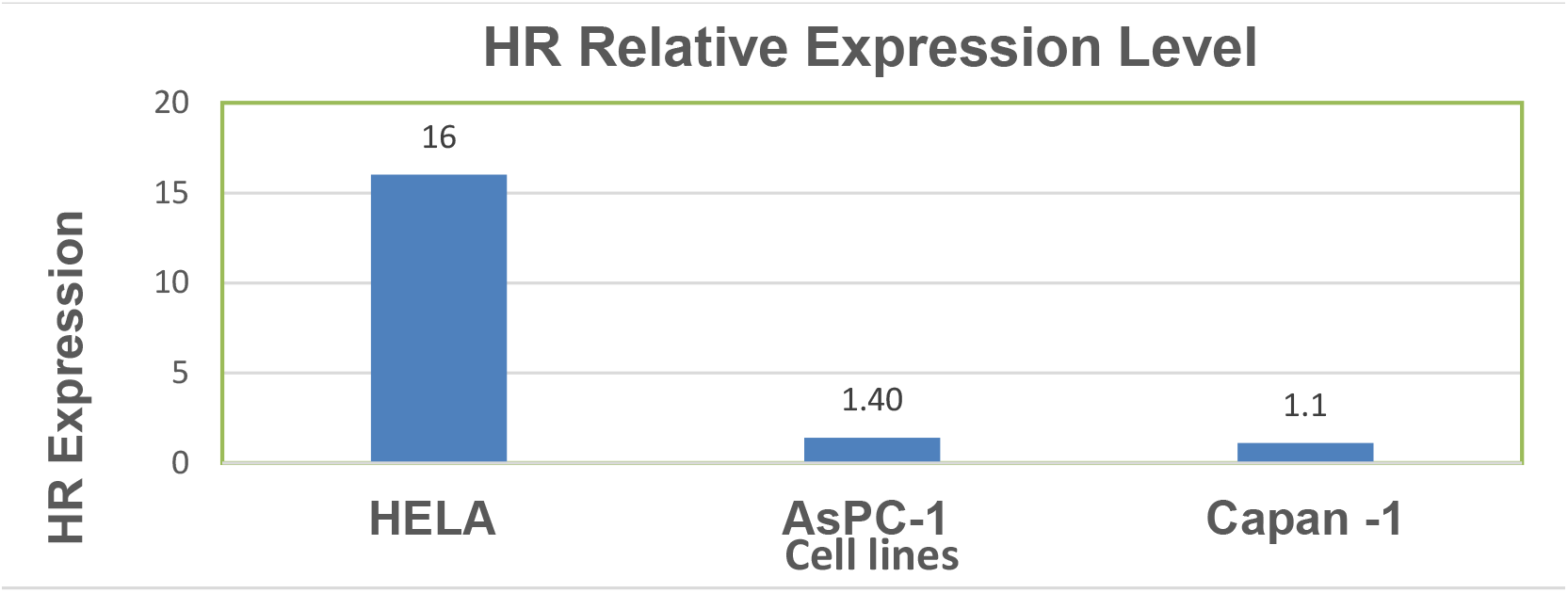
Real-time PCR results showing HR expression level in different cells line. The 16-fold HR expression level is found in HeLa Cell line whilst 1.4-fold and 1.1-fold HR expression is found AsPC-1 and CAPAN-1 cell lines, respectively. HR expression level reflects the level of HR repair pathway efficiency in that cell lines.

Although gel electrophoresis image results evidenced the presence of recombinant product following transfection (fig.11&12), the AsPC-1 and CAPAN-1 amplification plot results reveal very low starting material (recombinant product) as the fluorescent signals were detected beyond cycle 30. Since the cycle number at which fluorescent signal is detected is inversely proportional to the quantity of recombinant product, the amplification plot result in figure 13 suggests a very low quantity of recombinant products in both AsPC-1 and CAPAN-1 cell lines (Norgen Biotek Coperation 2010; Caraguel et al. 2011). This is further demonstrated by the analysis of Cq data shown in figure 14 below. The extracted Cq data were calculated to determine HR expression level, which is found to be 1.4 and 1.1-fold in AsPC-1 cells and CAPAN-1 cell line, respectively.

**Figure 12:**
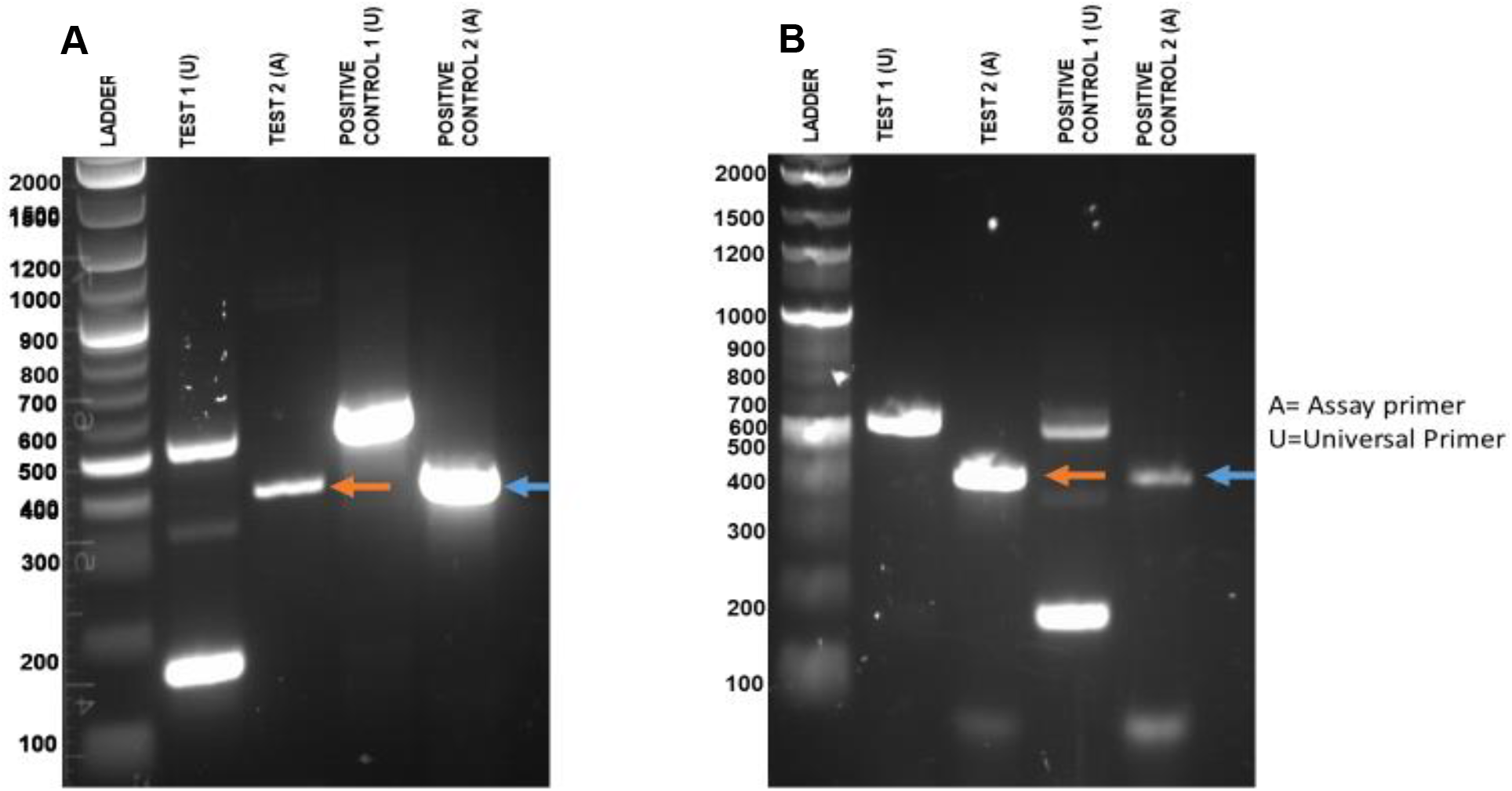
First and second transfection gel image results of CAPAN-1 cell line. A and B) the pink arrow indicates the homologous recombinant band (420bp) from both first and second transfections in test sample 2 and the blue arrow indicates the presence of homologous recombinant product (420bp) in the positive control as well. (Test 1(U) = test sample with universal primer; Test 2(A)= test sample 2 with assay primer; positive control 1(U)=positive control with universal primer; positive control 2(A)= positive control with assay primer; A= First transfection gel image; B= second transfection gel image; ladder=1kb plus)

## DISCUSSION

HR pathway is important during normal cellular proliferation and growth for accurate repair of DSBs. Due to such significant role of ensuring accurate repair of DSBs, HRD have been identified in cancers ranging from malignant tumours (colorectal, breast and ovarian cancers) to haematological malignancies such as AML (Cerbinskaite et al. 2012; Gaymes et al. 2002; Gaymes et al. 2013; Cream et al. 2017). The recent study of Heeke et al. (2018) have reported the prevalence of HRD across 21 solid tumours to be around 17.4%, highlighting the association of HRD with wide spectrum of cancers. Cancers such as breast and ovarian cancers with HRD exhibited hypersensitivity to DNA repair inhibitors such as PARPi because of synthetic lethality effects (Nickollof et al. 2017). Clinical stratification of newly diagnosed cancer patients defined based upon HR efficiency status is essential as cancer patients with HRD would benefit from PARPi, especially in combination with chemotherapy drugs such as Cisplatin. As evident in the case of breast and ovarian cancers, the combinatorial treatment approach resulted to improved survival. Such stratification would be a significant step toward precision medicine, treatment based upon tumour-biology (Brynat et al. 2005; Farmer et al. 2005; Drew and Plummer 2010; Audeh et al. 2010; Graeser et al. 2010).

The study aimed to optimize Norgen’s HR kit that can potentially be employed to quantify HR efficiency in cells. HR efficiency was quantified in both HR-proficient cancer cell lines (HeLa and AsPC-1) as well as an established HR-deficient cell line, CAPAN-1 cell line (Holt et al. 2008). The detection of the recombinant product in the test sample indicates successful HR between the two plasmids (dl-1 and dl-2) that led to generation of functional *LacZα* gene in HeLa cell line (fig.10A). Also, the real-time PCR showed high quantity of recombinant product generated in HeLa cells (figure.13), which is evidenced in the further analysis showing 16-fold increase in HR expression level in HeLa cell line (fig.14). Generally, the detection of recombinant product before cycle 30 of amplification curve is suggestive of high quantity of target sequence. The finding of elevated HR efficiency in HeLa cells from our study is in concordance with the previous studies (Yuan et al. 2012; Kim et al. 2018). For instance, the study of Yuan et al. (2012) showed that HeLa cell line exhibited greater than 8-fold increase in HR proficiency compared to HR-deficient cell lines with dysfunctional BRCA2 gene (Capan-1, and HCC1599) and BRCA1 gene (MDA-MBA436), when zinc finger nuclease assay was used (Mio et al. 2011). Melt curve rules out the presence of contaminant as single, sharp peaks were shown, suggesting that single, specific PCR product was generated with these sets of universal and assay primers (Valasek and Repa 2005).

We next repeated the procedure with another HR-proficient cell line, AsPC-1, to verify whether the kit would be able to quantify HR efficiency. Unfortunately, the real-time PCR results reveal low quantity of recombinant product in AsPC-1 cell line, reflecting overly decreased HR efficiency level (1.4-fold HR expression) when compared to 16-fold HR expression in HeLa cell line (Fig.13&14). Such level of decreased HR efficiency was not expected in AsPC-1 owing to its HR-proficient characteristic, allowing us to question the suitability of the PCR-based HR assay kit to quantify HR efficiency in other cell lines. However, the low HR efficiency found in AsPC-1 cell line might have arisen due to lost-of-functional mutations in some of the HR repair pathway genes such as Rad51, BRCA1/2 and MRN complex; this would interfere with the efficiency of HR process in AsPC-1 cell line (Cerbinskaite et al. 2012). Thus, further tests with the well-known established assays for HR efficiency such as immunofluorescence-based technique and gene expression study should be performed to confirm the HR repair pathway efficiency in the AsPC-1 cell line (Gaymes et al. 2009; Cerbinskaite et al. 2012).

CAPAN-1 cell line remains an important cell line in HR repair pathway studies as the pancreatic carcinoma cell line is known for its established HRD status. CAPAN-1 cell line is HR deficient owing to germline BRCA2 (617delT) mutation which cause a premature C-terminal truncation inactivating the domain for binding to Rad51. The C-terminal truncation of BRCA2 also inhibit the translocation of BRCA2 protein into the nucleus in response to the DSB due to missing C-terminal nuclear localization signals. BRCA2 protein has a downstream effect in HR repair pathway by mediating the recruitment of Rad51, which initiates homologous template search and then strand invasion following displacement of ssDNA stabilizer, RPA (Holt et al. 2008; Cerbinskaite et al. 2012; Hanamshet et al. 2016). The real-time PCR results demonstrated low quantity of recombinant product generated in CAPAN-1 cells, reflecting overly decreased HR efficiency level (1.4-fold HR expression) when compared to 16-fold HR expression in HeLa cell line (fig.13&14). This result re-affirms the specificity of PCR-based HR kit in detecting low level of HR efficiency in HRD cell line.

In conclusion, our results showed high quantity of recombinant product generated in HeLa cell line and low quantity in CAPAN-1 and AsPC-1 cell line which is interpreted as high HR efficiency in HeLa cell line whilst low HR efficiency in CAPAN-1 and AsPC-1 cell lines. Overall, the optimisation study has provided some evidence that the kit is appropriate enough for the quantification of HR pathway efficiency provided appropriate transfection reagent is used. Further study is required to confirm HR pathway efficiency in AsPC-1 cell line as the PCR-based kit revealed an overly decreased HR efficiency in this HR-proficient cells. Moreover, our optimization study has shown that the recommended protocol for real-time PCR condition is enough to accurately quantify the quantity of recombinant product generated in a successfully transfected cell line.

## Notes

### Competing Interest Statement

The authors have declared no competing interest.

